# A phosphorylation-dependent mechanism controls splice variant-specific S-palmitoylation of cardiac Kv4.3

**DOI:** 10.64898/2026.07.03.735752

**Authors:** Eleanor Dickson-Murray, Chunyun Du, Chien-Wen S Kuo, Rebecca J Gilchrist, Sheon Mary, Samuel Adu, Outi Kämäräinen, Amr Wahdan, Emma Dunning, Fiona Jordan, Elaine Brown, Jack W Houghton, Daria Berezina, Jianan Lu, Edward W Tate, Rachel C Myles, Francis L Burton, Godfrey L Smith, Helen Walden, Jules C Hancox, William Fuller

## Abstract

S-palmitoylation modulates the activity of many cardiac ion channels, yet the upstream signals that control this post-translational modification (PTM) are poorly defined. Here we identify a phosphorylation-dependent mechanism that governs splice variant-specific S-palmitoylation of the potassium channel Kv4.3. Using Acyl-RAC in native tissue and heterologous cells, we map palmitoylation to C546/547 in the intrinsically disordered Kv4.3 C-terminal tail. The short Kv4.3 splice variant Kv4.3S is ∼2.5-3-fold more palmitoylated than the long splice variant Kv4.3L, and systematic mutagenesis localises the dominant inhibitory determinant of Kv4.3 S-palmitoylation to residues 488-498 within the Kv4.3L-specific splice insert. KChIP2.1 promotes accumulation of a post-translationally modified Kv4.3 species that is selectively S-palmitoylated, and nanobody-targeted dephosphorylation removes this species and reduces Kv4.3 S-palmitoylation. Phos-tag electrophoresis and C-terminal truncation mapping identify S538 as the principal phosphorylation site enabling Kv4.3 S-palmitoylation; mutation of S538 markedly reduces formation of the palmitoylation-competent species. TurboID proximity-labelling and biochemical assays indicate that phosphorylation enhances recruitment of Kv4.3 to zDHHC5. Acute kinase inhibition rapidly eliminates phosphorylation but only gradually reduces palmitoylation, revealing temporal uncoupling between these PTMs. Functionally, non-palmitoylatable Kv4.3S exhibits larger peak currents, faster inactivation, and a left-shifted activation curve, consistent with palmitoylation limiting channel function and modulating gating transitions. Together, these findings identify phosphorylation of S538 as a priming modification that licenses Kv4.3 S-palmitoylation at C546/547, explain splice-variant differences in S-palmitoylation, and define a PTM cascade that tunes Kv4.3 channel behaviour.

## Introduction

Coordinated cardiac electrical activity is essential for normal cardiac function; electrical instability is a hallmark of many cardiac pathologies.^1, 2^ The cardiac action potential (AP) reflects the tightly controlled activity of inward depolarising and outward repolarising currents across the cardiomyocyte plasma membrane.^3–6^ Cardiac potassium channels, responsible for regulating cardiac repolarisation and resting membrane potential, are the most functionally diverse family of cardiac ion channels. Broadly, they can be divided into three categories: voltage-gated, inward rectifying and background potassium currents.^5, 7^ Over 40 different voltage-sensitive (K_v_) channels are expressed across multiple tissues in both excitable and non-excitable cells. Generally, these K_v_ channels consist of four pore forming α-subunits and multiple β-subunits with a range of interacting accessory subunits that fine tune channel function. K_v_ channels can either be homo-oligomers or hetero-oligomers.^5, 8^

During phase 1 of the cardiac AP the transient outward current (I_to_) transiently repolarises ventricular myocytes, regulating AP plateau voltage, morphology and duration. I_to_ is composed of two currents, I_to1_ and I_to2_. I_to1_ carries a potassium current that activates and inactivates rapidly and can be divided into fast (I_tof_, formed by Kv4.2/4.3) and slow (I_tos_, formed by Kv1.4/1.7/3.4).^9, 10^ Kv4.3 is a member of the Kv4 (Shal-related) family that mediates outward A-type potassium currents.^5, 8, 11^ The expression of I_tof_ and I_tos_ differs across species, giving rise to the different AP waveforms.^12–14^

Kv4.3 contains four pore forming α-subunits, with each subunit containing six transmembrane domains (S1-S6) and intracellular N and C termini. The N-terminal tetramerization (T1) domain is required for assembly of the four α-subunits.^15–17^ The intracellular C-terminal tail of Kv4.3 is largely unstructured.^16, 18^ Various accessory proteins interact with and regulate the activity of Kv4.3, including the KCNEs, DPPs, SAP97, and KChIPs.^19–26^ For example, KChIPs interact with both termini of Kv4.3 to stabilise the complex.^16^ Co-expression with KChIP2 significantly increases Kv4.3 cell surface expression, slows I_to_ activation and enhances I_to_ current amplitude.^11, 16, 24, 27–30^ KChIP2 expression is downregulated in heart failure (HF) and this has been suggested to contribute to changes in both calcium handling and repolarisation.^31^

Unique to Kv4.3 within the Kv4 family is the presence of an additional exon which is included in ∼50% of transcripts in the heart. Compared to the short variant Kv4.3S and other Kv4 family members, the alternatively spliced long variant Kv4.3L gains a 19 amino acid region (resides 488-506) within the S6 proximal unstructured cytosolic C-tail.^32, 33^ The current density, gating kinetics and voltage dependence of homomeric Kv4.3L and Kv4.3S recorded in murine fibroblasts at room temperature are indistinguishable.^34^ However, Kv4.3L contains a consensus phosphorylation site, T504, which is phosphorylated by protein kinase C (PKC). PKC both reduces peak Kv4.3-mediated currents in mammalian cells,^34^ and differentially modulates closed-state inactivation of Kv4.3L in *Xenopus* oocytes.^35^ Expression of the spliced forms remodels differently during the evolution of heart failure. Kv4.3L expression is upregulated ∼33% and Kv4.3S is downregulated ∼75% in the failing heart. This highlights the need for a greater understanding of the splice variants as potential therapeutic targets in heart failure.^36^

Lipid modifications, particularly those involving the attachment of a fatty acid, are the most common type of post-translational modification (PTM). These lipid modifications contribute to the stereochemical diversity of proteins and can be reversible or irreversible. S-palmitoylation, sometimes referred to as S-acylation or palmitoylation, refers to the reversible attachment of a C16 fatty acid, typically palmitic acid, to a cysteine residue via a thioester bond. S-palmitoylation reversibly anchors intracellular regions of proteins to membranes and can alter protein trafficking, protein-protein interactions and protein activity.^37–39^

In cardiomyocytes, multiple ion channels and their regulatory proteins are S-palmitoylated including HCN4, Nav1.5, the α1C-subunit of Cav1.2, NCX, KChIP2.1 and phospholemman.^40–47^ Here, we show that Kv4.3 is S-palmitoylated at C546/547 within its cytoplasmic C-terminal tail and that the Kv4.3S splice variant is more heavily palmitoylated than Kv4.3L. By systematic mutagenesis, we identify residues within the N-terminal half of the 19-amino-acid splice insert as primary determinants of this difference. We further demonstrate that phosphorylation promotes acquisition of a palmitoylation-competent Kv4.3 species, consistent with a model in which site-specific phosphorylation primes the Kv4.3 C-tail for S-palmitoylation at C546/547. Functionally, we show that loss of palmitoylation produces a gain-of-function phenotype characterised by increased current amplitude and altered gating kinetics, indicating that S-palmitoylation acts as a negative regulator of Kv4.3 activity. Together, these findings define a phosphorylation-dependent mechanism and cis-sequence elements that govern splice-variant-specific palmitoylation of Kv4.3.

## Results

### Kv4.3 is S-palmitoylated at C546/547

Kv4.3 and its homologues have been identified in several mammalian palmitoyl proteomes,^48–51^ but to date no palmitoylation sites have been identified. We used acyl-resin assisted capture (acyl-RAC) to assess Kv4.3 palmitoylation in mouse heart and brain tissue and in HEK293 cells transiently expressing Kv4.3L. Across all contexts, Kv4.3 was robustly recovered in Acyl-RAC eluates, indicating S-palmitoylation under native and heterologous conditions (Figure 1A). Systematic cysteine substitution in HEK293 cells mapped palmitoylation to C546/547 within the Kv4.3 intrinsically disordered C-terminal tail. Deleting the N-terminal T1 domain also substantially reduced palmitoylation of full-length Kv4.3L, placing channel assembly upstream of a palmitoylation-competent state (Figure 1B). This T1 dependence of Kv4.3 palmitoylation is consistent with the idea that proper tetramerization and/or a specific inter-domain juxtaposition is required to expose C546/547 to the acyltransferase active site.^17, 52^

**Figure 1:**
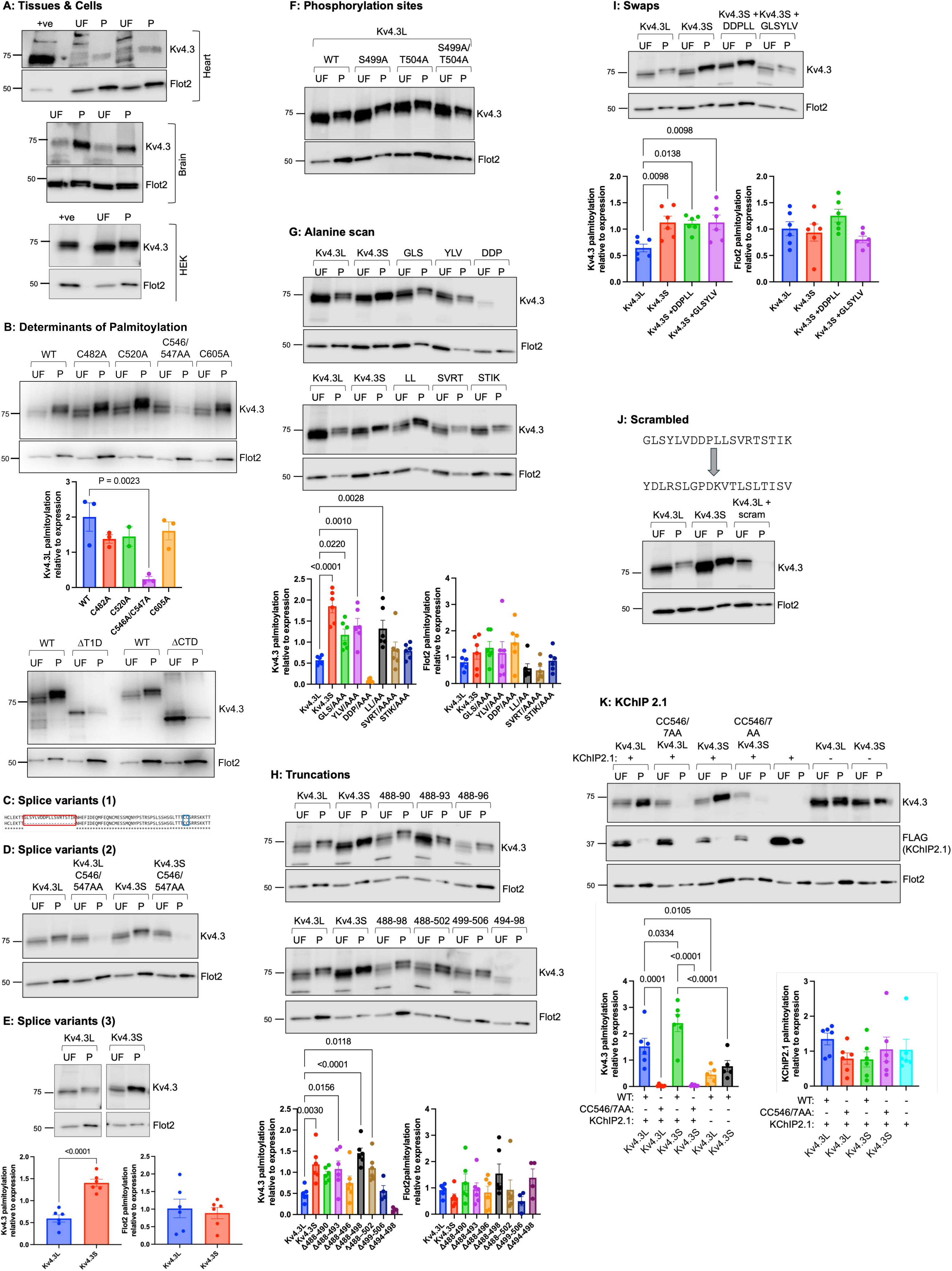
Sequence, domain, and subunit influence on Kv4.3 palmitoylation. **A:** Kv4.3 is palmitoylated. Acyl-RAC analysis of heart and brain tissue from female (left) and male (right) mice and HEK293 cells transfected with Kv4.3L. Samples were immunoblotted for Flot2 as a control for the Acyl-RAC experiment. UF: unfractionated lysate; P: purified palmitoylated proteins. **B:** Determinants of Kv4.3 palmitoylation. Acyl-RAC analysis of HEK293 cells transfected with wild type Kv4.3L or a series of Kv4.3L point mutations and deletions. WT: wild type, ΔT1D: T1 domain deleted, ΔCTD: intracellular C tail deleted. The graph shows mean±SEM palmitoylation relative to expression for each mutant, N=2-3 per group, analysed with a one-way ANOVA with a Dunnett’s post hoc test. **C:** Kv4.3 splice site. Sequence alignment of Kv4.3L 481-555 with Kv4.3S 481-536. Red box indicates the 19 amino acid splice region, and the blue box indicates S-palmitoylation sites C546/547 (long variant numbering). **D:** Palmitoylation of Kv4.3 splice variants. Acyl-RAC analysis of HEK293 cells transfected with either WT Kv4.3L, Kv4.3S, or C546/547AA (long variant numbering) confirms the same cysteines are palmitoylated in both variants. **E:** Kv4.3S is more palmitoylated than Kv4.3L. Acyl-RAC analysis of HEK293 cells transfected with either WT Kv4.3L or Kv4.3S. The graph shows mean±SEM palmitoylation relative to expression for each Kv4.3 variant or Flot2, N=6 per group, analysed with an unpaired t-test. **F:** Kv4.3L-specific phosphorylation does not explain differential palmitoylation. Acyl-RAC analysis of HEK293 cells transfected with either WT Kv4.3L or alanine mutants of established phosphorylation sites unique to Kv4.3L. **G:** Alanine scanning mutagenesis of Kv4.3L. Acyl-RAC analysis of HEK293 cells transfected with either WT Kv4.3L or alanine mutants of the splice site. GLS: G488A/L489A/S490A, YLV: Y491A/L492A/V493A, DDP: D494A/D495A/P496A, LL: L497A/L498A, SVRT: S499A/V500A/R501A/T502A, STIK: S503A/T504A/I505A/K506A. The graphs show mean±SEM palmitoylation relative to expression for each Kv4.3L mutant or Flot2, N=6 per group, analysed with a one-way ANOVA with a Dunnett’s post hoc test. **H:** Truncation mutagenesis of Kv4.3L. Acyl-RAC analysis of HEK293 cells transfected with either WT Kv4.3L or truncation mutants of the splice site. The graphs show mean±SEM palmitoylation relative to expression for each Kv4.3L mutant or Flot2, N=6 per group, analysed with a one-way ANOVA with a Dunnett’s post hoc test. **I:** Insertional mutagenesis of Kv4.3S. Acyl-RAC analysis of HEK293 cells transfected with either WT Kv4.3S or Kv4.3S into which selected regions of the N terminal half of the Kv4.3L splice site are inserted. The graphs show mean±SEM palmitoylation relative to expression for each Kv4.3 variant or Flot2, N=6 per group, analysed with a one-way ANOVA with a Dunnett’s post hoc test. **J:** Scrambling the splice site in Kv4.3L. Acyl-RAC analysis of HEK293 cells transfected with WT Kv4.3L, WT Kv4.3S, or Kv4.3L in which the splice site sequence is scrambled. **K:** Impact of co-expressing KChIP2.1 on Kv4.3 palmitoylation. Acyl-RAC analysis of HEK293 cells transfected with either WT Kv4.3S or WT Kv4.3L with or without KChIP2.1. The graphs show mean±SEM palmitoylation relative to expression for each Kv4.3 variant or KChIP2.1, N=5-6 per group, analysed with a one-way ANOVA with Šidák’s post hoc test.

### Differential S-palmitoylation of Kv4.3 splice variants

Unique to Kv4.3 within the Kv4 family is the presence of an additional spliced region. The alternatively spliced version, Kv4.3S, lacks a 19 amino acid region (residues 488-506, Kv4.3L numbering) within the S6 proximal cytosolic C-terminus compared to Kv4.3L (Figure 1C). We first confirmed that the same cysteines are palmitoylated in both splice variants (Figure 1D). Notably, Kv4.3S was significantly more S-palmitoylated than Kv4.3L (Figure 1E). Because phosphorylation and S-palmitoylation can antagonise each other, we first tested whether phosphorylation within the 19-residue spliced segment explains the difference. Mutations of two established phosphorylation sites in the alternatively spliced region, T504A or S499A, did not alter the S-palmitoylation of Kv4.3L to match that of Kv4.3S (Figure 1F).^19, 35^ We conclude from these experiments that Kv4.3L is not less palmitoylated than Kv4.3S because of phosphorylation in the alternatively spliced region.

To define sequence determinants of differential splice variant palmitoylation, we performed complementary perturbations: (i) alanine scanning across residues 488-506, (ii) stepwise truncations, (iii) insertion of sub-segments into Kv4.3S, and (iv) complete scrambling of the 19-residue splice site. All mutants were expressed in HEK293 cells, and their S-palmitoylation assessed via Acyl-RAC (Figure 1G). Alanine substitutions at GLS (488-490), YLV (491-493), and LL (497-498) but not SVRT (499-502) or STIK (503-506) each increased Kv4.3L palmitoylation. Alanine mutants of the DDP block (494-496), which matches a DXXLL endosomal targeting motif^53^ expressed poorly. Kv4.3L truncations removing residues 488-493, 488-498, or 488-502 (but not 499-506) likewise enhanced palmitoylation (Figure 1H), localising the strongest negative influence on Kv4.3L palmitoylation to the N-terminal half of the splice site (488-498). Truncation mutants of the DDPLL block (494-498) expressed poorly. Introducing only GLSYLV (488-493) or DDPLL (494-498) into Kv4.3S did not depress its palmitoylation (Figure 1I). A scrambled-sequence Kv4.3L variant expressed but was not detectably palmitoylated and accumulated to lower abundance than wild-type (Figure 1J). Together, these experiments indicate that residues 488-498 (GLSYLVDDPLL) act as a cis-determinant that reduces palmitoylation of Kv4.3L, with GLS, YLV, and LL contributing most.

### KChIP2.1 enhances Kv4.3 palmitoylation without abolishing splice-variant differences

Using a bicistronic Kv4.3-IRES-KChIP2.1 vector, we asked whether the auxiliary subunit modifies Kv4.3 palmitoylation. Co-expression with KChIP2.1 significantly increased palmitoylation relative to expression of both Kv4.3 splice variants alone, yet Kv4.3S remained more palmitoylated than Kv4.3L (Figure 1K). KChIP2.1 itself was palmitoylated but its palmitoylation was unchanged by co-expression with either Kv4.3 variant (Figure 1K), indicating that KChIP2.1 promotes a palmitoylation-competent conformation / cellular location rather than being reciprocally modified by the pore-forming subunit.

### A slower-migrating, palmitoylation-competent Kv4.3 species requires KChIP2.1 and N-terminal architecture

To test whether a specific Kv4.3 conformation is prerequisite for palmitoylation of the distal C-tail, we engineered a transmembrane-deleted mutant (TMD-del) containing the N-terminus, the linker to the first transmembrane domain, and the full C-terminus (Figure 2A). TMD-del was not palmitoylated when expressed alone. Strikingly, co-expression with KChIP2.1 restored palmitoylation of TMD-del and yielded two Kv4.3 bands on SDS-PAGE; only the slower migrating species was enriched by Acyl-RAC (Figure 2B). The slower band was also observed for both TMD-del splice variants and a non-palmitoylatable mutant co-expressed with KChIP2.1 (Figure 2C). Replacing KChIP2.1 with KChIP2.2, or deleting the N-terminus or T1 region from Kv4.3, abolished the slower band (Figure 2D). Thus, KChIP2.1 and the Kv4.3 N-terminal/T1 architecture are necessary for formation of a slower-migrating species that is competent for palmitoylation, independent of splice form or cysteine palmitoylation state.

**Figure 2:**
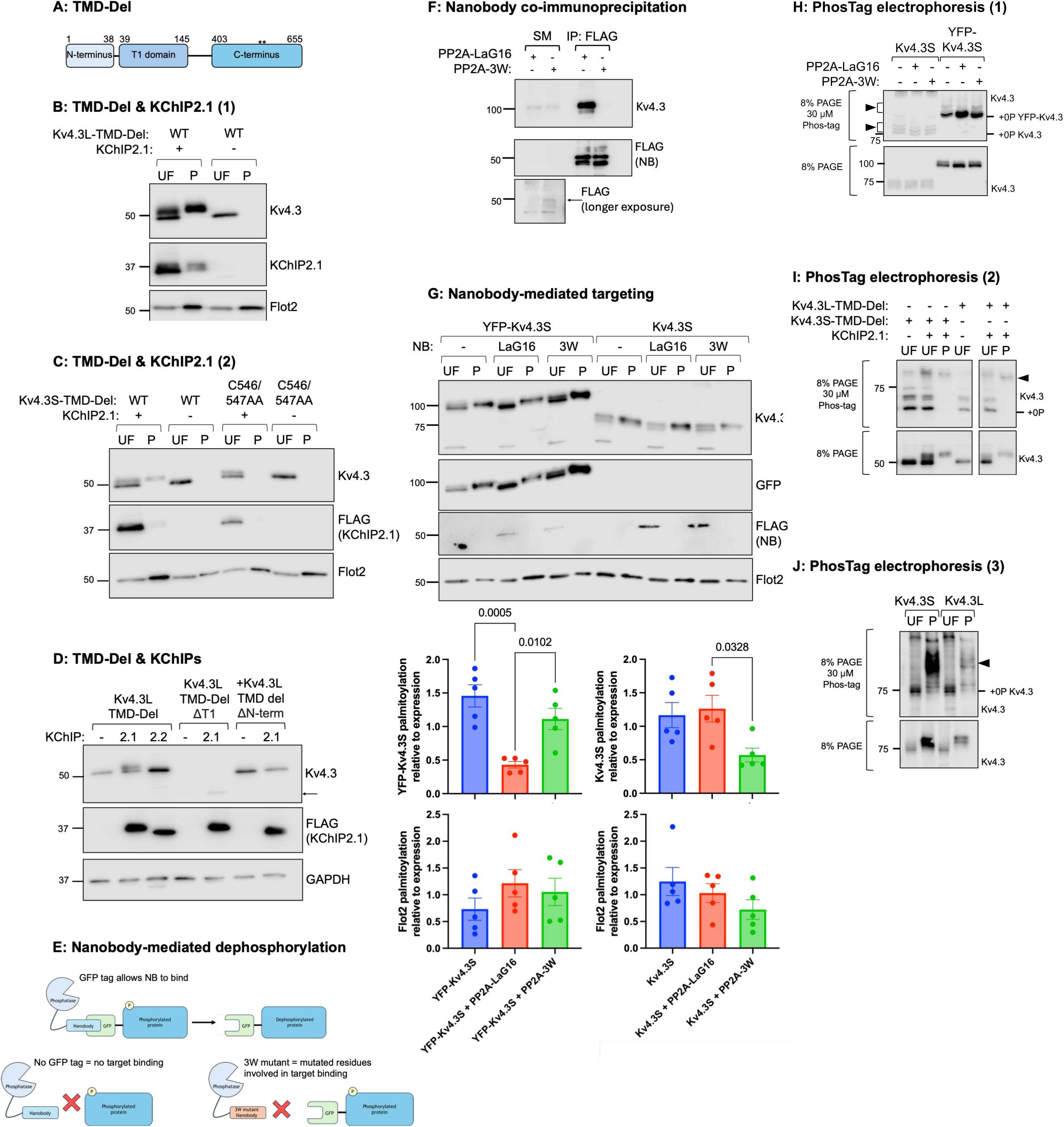
Kv4.3 phosphorylation promotes its palmitoylation. A: TMD-Del schematic. TMD-Del is composed of N terminal, T1, and C terminal regions of Kv4.3 (long variant numbering). The positions of the palmitoylation sites are indicated by **. **B:** Palmitoylation of TMD-Del. Acyl-RAC analysis of HEK293 cells transfected with TMD-Del Kv4.3L with and without KChIP2.1. Samples were immunoblotted for Flot2 as a control for the Acyl-RAC experiment. UF: unfractionated lysate; P: purified palmitoylated proteins. **C:** KChIP2.1 induced bandshift is independent of palmitoylation. Acyl-RAC analysis of HEK293 cells transfected with TMD-Del Kv4.3S (WT or non-palmitoylated) with or without KChIP2.1. **D:** TMD-Del bandshift requires Kv4.3 N terminal domains and KChIP2.1. Whole cell lysates from HEK293 cells expressing TMD-Del Kv4.3L and truncation mutants with or without KChIP2.1 or KChIP2.2. ΔN-term: residues 1-38 deleted, ΔT1: residues 39-145 deleted. The arrow indicates the position of the TMD-del ΔT1 protein. **E:** Nanobody schematic. Nanobody (NB) binding with the GFP tag brings PP2A into proximity with the target protein. The nanobody is unable to deliver PP2A to proteins not tagged with GFP. The 3W nanobody has had one amino acid in each complimentary determining region mutated to a tryptophan (F30W/T55W/G103W) to reduce its affinity for GFP and prevent delivery of PP2A. **F:** Confirmation of nanobody/target interaction. FLAG tagged nanobody-PP2A chimeras were co-expressed with YFP-Kv4.3 in HEK293 cells and immunoprecipitated. SM: soluble lysate, IP: immunoprecipitation fraction. The arrow indicates the position of PP2A-NB. **G:** Nanobody-mediated targeting of Kv4.3S palmitoylation. Acyl-RAC analysis of HEK293 cells transfected with YFP-Kv4.3S or Kv4.3S with either no nanobody, PP2A-LaG16, or PP2A-3W. The graphs show mean±SEM palmitoylation relative to expression for YFP-Kv4.3, Kv4.3 or Flot2, N=5 per group, analysed with a one-way ANOVA with Tukey’s post hoc test. **H:** Kv4.3 dephosphorylation. Whole cell lysates from HEK293 cells expressing Kv4.3S or YFP-Kv4.3S with either no nanobody, PP2A-LaG16, or PP2A-3W analysed using Phos-Tag PAGE (upper) and standard SDS PAGE (lower). Slower migrating phosphorylated forms of YFP-Kv4.3 but not Kv4.3S (arrowheads) are removed only by PP2A-LaG16. **I:** TMD-Del phosphorylation and palmitoylation. Whole cell lysates (UF) or purified palmitoylated proteins (P) from HEK293 cells expressing TMD-Del-Kv4.3S or TMD-Del-Kv4.3L with or without KChIP2.1 analysed using Phos-Tag PAGE (upper) and standard SDS PAGE (lower). The presence of slower migrating phosphorylated forms of Kv4.3 (arrowheads) is promoted by co-expressing KChIP2.1. Only these phosphorylated forms are palmitoylated. **J:** Kv4.3 phosphorylation and palmitoylation. Whole cell lysates (UF) or purified palmitoylated proteins (P) from HEK293 cells expressing Kv4.3S or Kv4.3L analysed using Phos-Tag PAGE (upper) and standard SDS PAGE (lower). Only slower migrating phosphorylated forms of Kv4.3 (arrowhead) are palmitoylated.

### Targeted dephosphorylation collapses the slower species and reduces palmitoylation

We hypothesised that the slower band represents a post-translationally modified Kv4.3 species that favours palmitoylation, focusing first on phosphorylation as the ‘priming’ modification. Nanobodies are single chain antibodies in which the VHH protein binding domain can fold in the mammalian cytoplasm to form a small stable domain that binds its target with high affinity. Many PTMs rely on protein-protein interactions: enzyme-nanobody chimeras are validated tools to deliver enzymes mediating PTMs to a nanobody’s target protein.^54^ We therefore fused the GFP/YFP/CFP binding nanobody LaG-16 to the protein phosphatase PP2A and co-expressed this reagent with YFP-Kv4.3. Control experiments utilised a nanobody in which one amino acid in each of the three complementarity-determining regions was mutated to tryptophan. This ‘3W’ form displays a reduced ability to bind and deliver enzymes to YFP (Figure 2E). LaG-16-PP2A co-immunoprecipitated with YFP-Kv4.3S but not with untagged Kv4.3 (Figure 2F). In cells expressing LaG-16-PP2A, YFP-Kv4.3S collapsed from a doublet to a single, faster band and showed reduced palmitoylation relative to controls (Figure 2G). Untagged Kv4.3S palmitoylation decreased modestly with PP2A-3W (but not with LaG-16-PP2A), possibly indicating some off-target activity. These findings support the concept that phosphorylation generates the slower migrating species and promotes palmitoylation.

We set out to directly visualise Kv4.3 phosphorylation using Phos-tag electrophoresis. The Phos-tag ligand retards the progress of phosphorylated proteins on SDS PAGE, allowing them to be visualised by a mobility shift. Phos-tag SDS-PAGE resolved three Kv4.3 mobility states, consistent with multiple phosphorylation states. LaG-16-PP2A shifted YFP-Kv4.3S toward the unphosphorylated state relative to the two controls (Figure 2H). TMD-del truncations of both splice variants were more phosphorylated in the presence of KChIP2.1 (Figure 2I), and Acyl-RAC-enriched Kv4.3 fractions were selectively phosphorylated (Figure 2J). Taken together, these results suggest that the TMD del mutant becomes phosphorylated in the presence of KChIP2.1 and that the palmitoylated form of Kv4.3 is considerably more phosphorylated. The data are therefore consistent with the concept that phosphorylation is required for Kv4.3 palmitoylation.

### Phosphorylation site mapping in Kv4.3

We set out to identify the phosphorylation site in Kv4.3 that promotes its palmitoylation by further truncating the TMD-del mutant (Figure 3A). All mutants were co-expressed with KChIP2.1. Their interaction with KChIP2.1 was assessed by co-immunoprecipitation, and their phosphorylation by the presence of a slower migrating species on SDS PAGE. The phosphorylation-induced change in Kv4.3 mobility was lost when 130 amino acids were deleted but present when 100 amino acids were deleted from the Kv4.3 C tail, indicating the phosphorylation site(s) are within this region (Figure 3B). FLAG co-immunoprecipitation confirmed that both the Δ100 and Δ130 deletion mutants still bind KChIP2.1 (Figure 3C). Further deletions (Figure 3D) and mutagenesis of candidate serine / threonine residues in Kv4.3 to alanine (Figure 3E) identified S537 and S538 as the key residues responsible for altered Kv4.3 mobility on SDS PAGE. Alanine mutagenesis of only S538 was sufficient to substantially reduce Kv4.3 palmitoylation (Figure 3F), implying phosphorylation here is required for Kv4.3 palmitoylation. We therefore generated pseudophosphorylated (S to D) mutations of both S537 and S538 in TMD-Del Kv4.3 and evaluated whether the dependence of TMD-Del palmitoylation on co-expressing KChIP2.1 was abrogated by positioning a negative charge in position 537 or 538 (Figure 3G). Palmitoylation of TMD-Del remained KChIP2.1-dependent in these experiments and was impaired by mutation S538D. S to D mutagenesis of S537 but not S538 in full length Kv4.3 slightly enhanced its palmitoylation (Figure 3H), suggesting aspartate mutagenesis of S538 cannot fully substitute for phosphorylation in promoting Kv4.3 palmitoylation. Taken together, we conclude that phosphorylation of Kv4.3 S538 is required for palmitoylation of Kv4.3 and is promoted by the presence of KChIP2.1, but we do not rule out that phosphorylation at other sites may also enhance Kv4.3 palmitoylation.

**Figure 3:**
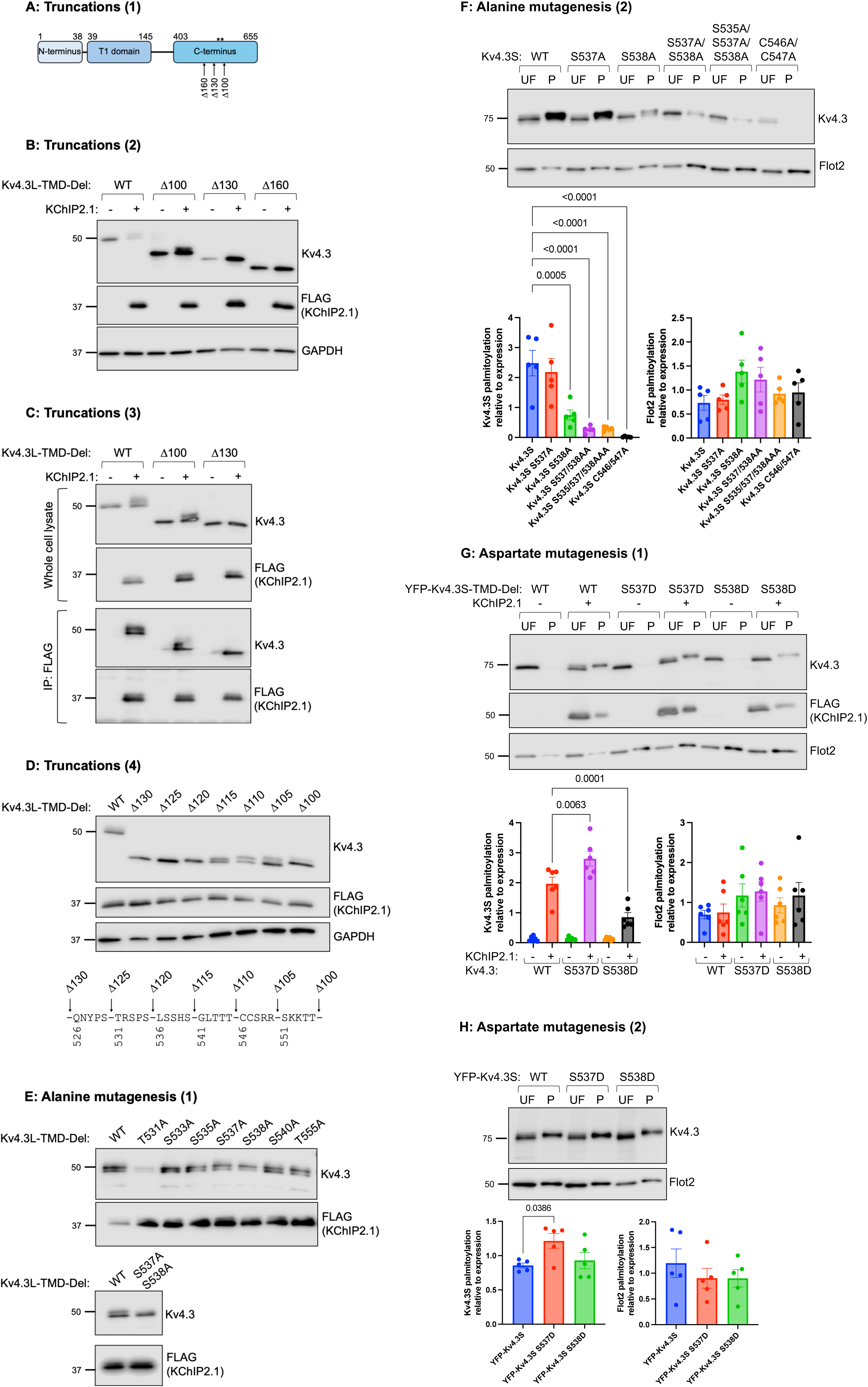
Phosphorylation site mapping in Kv4.3. A: TMD-Del truncation schematic. Truncation positions of mutants deleting 100, 130, 160 amino acids from the TMD-Del C terminus. The positions of the palmitoylation sites are indicated by **. **B:** Truncations to identify the Kv4.3 phosphorylation site. Whole cell lysates from HEK293 cells expressing TMD-del Kv4.3L truncations with and without KChIP2.1 analysed by SDS PAGE. The KChIP2.1-dependent slower migrating phosphorylated form of Kv4.3 is present in Δ100 but absent from Δ130 and Δ160. **C:** KChIP2.1 binding of TMD-del truncations. Immunoprecipitation (IP) of FLAG tagged KChIP2.1 confirms both Δ100 and Δ130 truncations of TMD-del Kv4.3L bind KChIP2.1. **D:** Fine mapping of the Kv4.3 phosphorylation site. Whole cell lysates from HEK293 cells expressing TMD-del Kv4.3L truncations and KChIP2.1 analysed by SDS PAGE. The KChIP2.1-induced change in mobility is lost when Kv4.3 536-LSSHS-540 is deleted. **E:** Candidate phosphorylation site mutagenesis. Whole cell lysates from HEK293 cells expressing KChIP2.1 and TMD-del Kv4.3L serine and threonine to alanine mutations analysed by SDS PAGE. No single mutation abolishes the KChIP2.1-induced change in mobility, but dual mutation of serines 537 & 538 does. **F:** Impact of phosphorylation site mutagenesis on Kv4.3 palmitoylation. Acyl-RAC analysis of HEK293 cells transfected with either WT Kv4.3S, alanine mutants of candidate phosphorylation sites, or non-palmitoylated C546/7AA Kv4.3. The graphs show mean±SEM palmitoylation relative to expression for each Kv4.3S mutant or Flot2, N=5 per group, analysed with a one-way ANOVA with a Dunnett’s post hoc test. **G:** Aspartate mutagenesis of candidate TMD-del Kv4.3 phosphorylation sites. Acyl-RAC analysis of HEK293 cells transfected with YFP-TMD-del Kv4.3S and aspartate mutants of candidate phosphorylation sites, with or without KChIP2.1. The graphs show mean±SEM palmitoylation relative to expression for each Kv4.3S mutant or Flot2, N=6 per group, analysed with a one-way ANOVA with a Šídák’s post hoc test. **H:** Aspartate mutagenesis of candidate Kv4.3 phosphorylation sites. Acyl-RAC analysis of HEK293 cells transfected with YFP-Kv4.3S and aspartate mutants of candidate phosphorylation sites. The graphs show mean±SEM palmitoylation relative to expression for each Kv4.3S mutant or Flot2, N=6 per group, analysed with a one-way ANOVA with a Tukey’s post hoc test.

### The relationship between Kv4.3 and zDHHC5

We next aimed to identify the zDHHC-PAT(s) responsible for palmitoylating Kv4.3 by fusing the proximity labelling enzyme TurboID to the Kv4.3 amino terminus and generating a cell line stably expressing TurboID-Kv4.3L. Proteins in proximity to Kv4.3 were labelled by briefly treating cells with biotin before lysis, and biotinylated proteins purified and identified using mass spectrometry. Among 691 proteins in proximity to Kv4.3 (Supplementary Table 1) were two zDHHC-PATs: the closely related enzymes zDHHC5 and zDHHC8. We investigated the relationship between zDHHC5 and Kv4.3. Reciprocal co-immunoprecipitation experiments identified a physical interaction between zDHHC5 and Kv4.3S (Figure 4A). Re-expression of zDHHC5 in a zDHHC5 knockout cell line increased Kv4.3S palmitoylation, suggesting a functional relationship between the two proteins (Figure 4B). Mutation of S537 and S538, the key phosphorylation site(s) required for Kv4.3 palmitoylation, reduced but did not abolish the physical interaction between Kv4.3S and zDHHC5 (Figure 4C), suggesting that phosphorylating Kv4.3 S538 promotes its recruitment by zDHHC5, but that other contact points exist between the two proteins.

**Figure 4:**
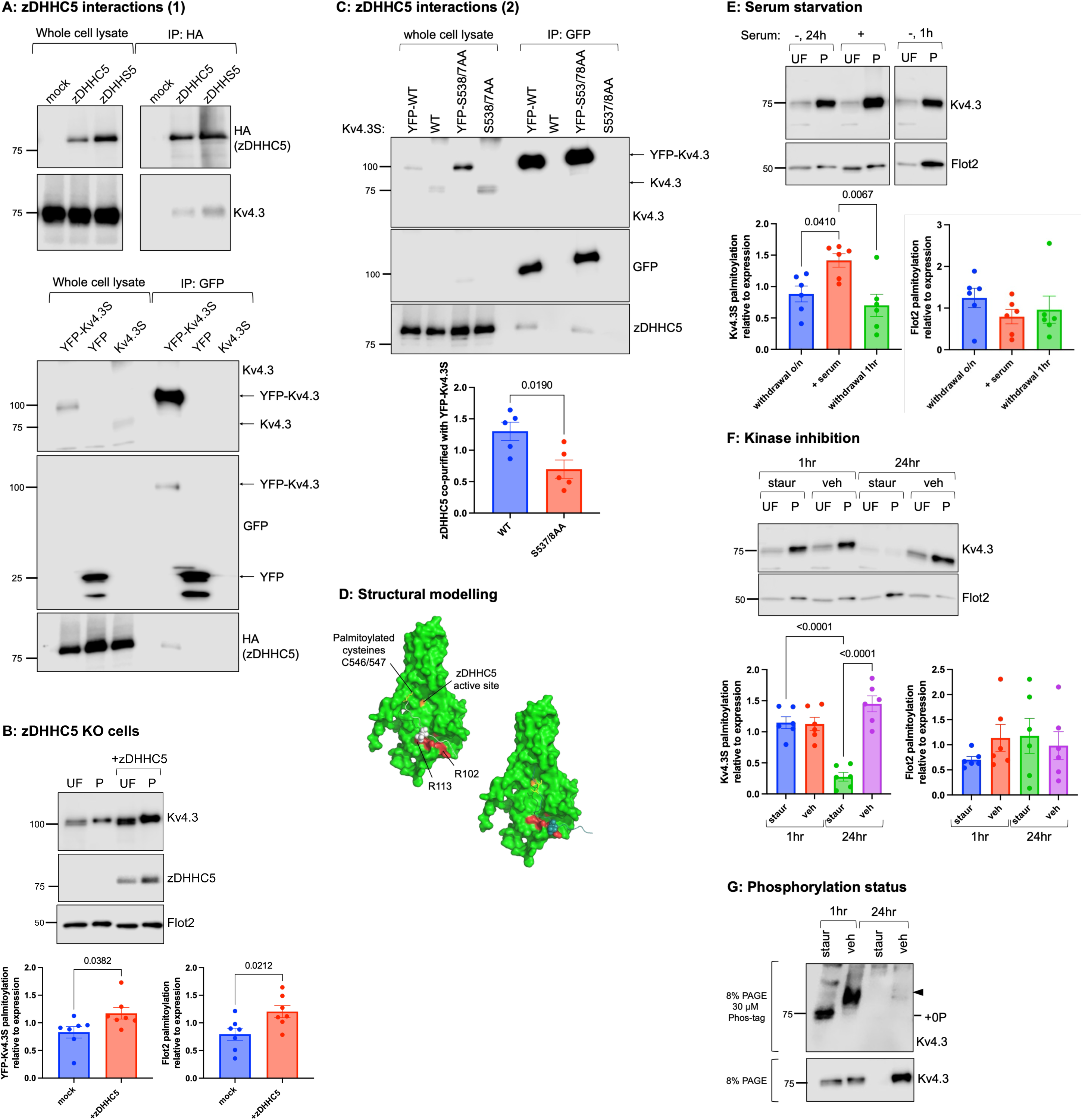
Phosphorylation primes Kv4.3 for palmitoylation by zDHHC5. A: A physical interaction between zDHHC5 and Kv4.3. Top: immunoprecipitation of HA-zDHHC5 and HA-zDHHS5 from FT293 cells stably expressing Kv4.3S. Bottom: immunoprecipitation of YFP-Kv4.3S (but not YFP alone) co-purifies HA-zDHHC5 from transiently transfected HEK293 cells. **B:** A functional interaction between zDHHC5 and Kv4.3. Acyl-RAC analysis of zDHHC5 KO cells transfected with WT YFP-Kv4.3S with or without zDHHC5. The graphs show mean±SEM palmitoylation relative to expression for YFP-Kv4.3S or Flot2, N=7 per group, analysed with an unpaired t-test. **C:** Impact of Kv4.3 phosphorylation on zDHHC5 interaction. Co-immunoprecipitation of endogenous zDHHC5 with YFP-Kv4.3S is reduced by mutation S537A/S538A. The graph shows the amount of zDHHC5 coimmunoprecipitated relative to the amount of YFP-Kv4.3S captured, N=5 per group, analysed with an unpaired t-test. **D:** Modelling zDHHC5 with Kv4.3 peptide. Model of zDHHC5 (AF-Q9C0B5-F1) with a phosphorylated Kv4.3 peptide (residues 531-550, including phosphoserine 538 shown with spheres and palmitoylation sites with yellow sticks) created using HPEPDOCK. The active site of zDHHC5 is orange and two positively charged amino acids on the zDHHC5 surface are coloured red. Two modelling scenarios in which the Kv4.3 palmitoylated cysteines closely approach the zDHHC5 active site are shown. **E:** Serum starvation and Kv4.3 palmitoylation. Acyl-RAC analysis of FT293 cells stably expressing Kv4.3S and serum starved for either 1 h or overnight. The graphs show mean±SEM palmitoylation relative to expression for Kv4.3S or Flot2, N=6 per group, analysed with an unpaired t-test. **F:** Kinase inhibition and Kv4.3 palmitoylation. Acyl-RAC analysis of FT293 cells stably expressing Kv4.3S and treated with either staurosporine (10 µM) or vehicle (0.1% DMSO) for either 1 h or overnight. The graphs show mean±SEM palmitoylation relative to expression for Kv4.3S or Flot2, N=6 per group, analysed with a one-way ANOVA with a Šidák’s post hoc test. **G:** Palmitoylation of unphosphorylated Kv4.3S. Purified palmitoylated proteins from FT293 cells stably expressing Kv4.3S treated with either staurosporine (10 µM) or vehicle (0.1% DMSO) for either 1 h or overnight analysed using Phos-Tag PAGE (upper) and standard SDS PAGE (lower). Treating with stauropsorine for 1 h generates a palmitoylated form of Kv4.3S which is not phosphorylated.

S-palmitoylation and phosphorylation are traditionally thought to oppose each other because they attract and repel proteins from membranes respectively. The AlphaFold predicted structure of zDHHC5 includes two positively charged pockets on the enzyme surface in the intracellular loop containing the catalytic site between transmembrane domains 2 and 3. This topology has recently been confirmed experimentally.^55^ We hypothesised that these regions could conceivably interact with and position a phosphorylated region of a protein to present it to the zDHHC5 active site. We therefore used HPEPDOCK to examine whether a phosphorylated Kv4.3 peptide (residues 531-550, containing the S-palmitoylation and phosphorylation sites) would be appropriately positioned for palmitoylation if phosphorylation promoted its interaction with zDHHC5. Models that did not position the palmitoylated cysteines close enough to the active site for catalysis were rejected. Several modelling scenarios suggest that phosphorylation of S538 favours Kv4.3 interaction with zDHHC5, and that there is enough flexibility in the 531-550 region of Kv4.3 to allow the S-palmitoylated cysteines C546/547 to be brought into proximity with the zDHHC5 active site (Figure 4D).

We used serum starvation and kinase inhibitors to identify the kinase family responsible for phosphorylating Kv4.3 S538. A significant decrease in Kv4.3S S-palmitoylation was observed following both an overnight and a one-hour serum withdrawal (Figure 4E), suggesting that a serum-dependent kinase may be responsible for S538 phosphorylation. A significant decrease in Kv4.3S S-palmitoylation was also observed when cells were treated with staurosporine overnight (Figure 4F), but treating cells with staurosporine for 1 hour caused no significant change in Kv4.3S S-palmitoylation. We examined the phosphorylation status of Kv4.3 using Phos-tag SDS-PAGE. This showed a phosphorylated species in vehicle treated cells which was entirely absent from cells treated with staurosporine for 1 hour, indicating that Kv4.3 is no longer phosphorylated but remains S-palmitoylated (Figure 4G). We conclude from these experiments that kinase inhibition for 1 hour quickly prevents Kv4.3 phosphorylation, however Kv4.3S remains S-palmitoylated until the phosphorylation has been prevented for a longer period, in this case overnight.

### Influence of palmitoylation on Kv4.3 function

We tested whether palmitoylation at C546/547 modulates Kv4.3 channel function by comparing WT Kv4.3S with a non palmitoylatable C546/547AA mutant in HEK293 cell lines stably expressing Kv4.3S using whole-cell patch clamp. Different protocols were applied to evaluate the electrophysiological kinetics of Kv4.3 channels.

The voltage dependence of activation was assessed using a standard protocol shown in the inset of Figure 5. Relative to wild type Kv4.3S, C546/547AA produced larger peak currents which inactivated more rapidly, with a left shifted V_0.5_ of activation and an increased slope factor (Figure 5Ai, 5Aii, 5B, 5C and 5D). The time-course of channel inactivation was evaluated by bi-exponential fitting of current elicited at +20 mV. The slow time constant of inactivation (τ_s_) was significantly smaller for C546/547AA than WT Kv4.3S, while the fast component of inactivation (τ_f_) was unaltered (Figure 5E).

**Figure 5:**
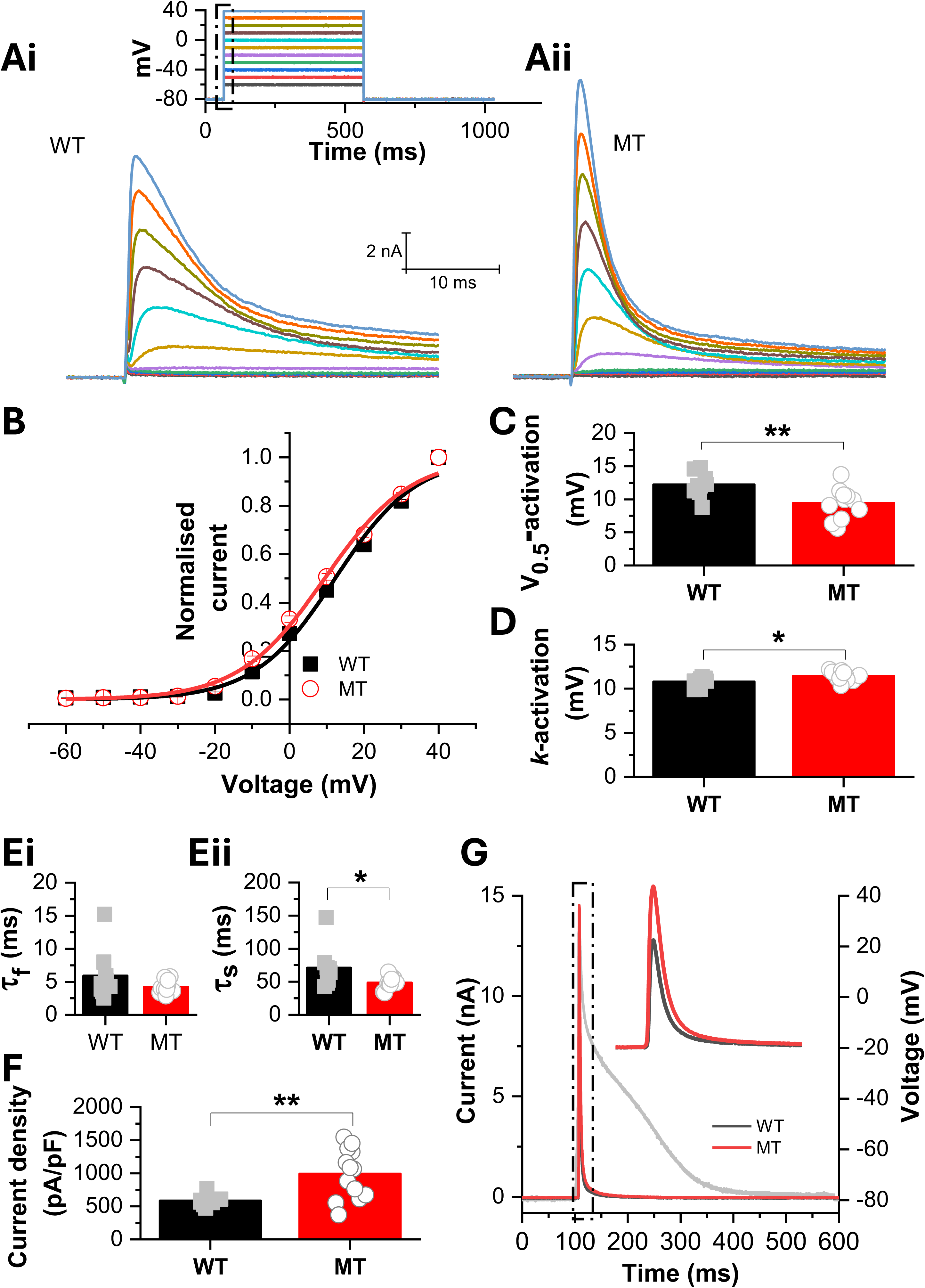
Effects of palmitoylation on Kv4.3S activation, time-course of inactivation and amplitude during atrial action potential waveform. A: Representative families of recordings from cells expressing wild type (WT, left, Ai) or non-palmitoylated C546/7AA (MT, right, Aii) Kv4.3S, elicited by the voltage protocol shown as an inset to ‘Ai’. The displayed traces focus on the current profile observed early following depolarisation (highlighted in the inset by the boxed area). Colour coding matches each voltage pulse to a current recording. **B:** Normalised current-voltage (I-V) plots for WT (black) and C546/547AA (red) Kv4.3S currents. For these, currents were normalised to maximal current during the protocol and the resulting mean plots were fitted with a Boltzmann equation to give half-maximal activation voltage (V_0.5_) and slope factor (*k*) values. **C** and D: Plots of mean ± SEM values for activation V_0.5_ (C, WT: 12.20±0.62 mV, C546/7AA: 9.40±0.66 mV) and *k* (D, WT: 10.77±0.18 mV, C546/7AA: 11.41±0.15 mV) respectively. Superimposed on the bar charts are the values obtained by fitting data from individual experiments (* P<0.05; **P<0.01, t-test; n=9 for WT and n=12 for C546/547AA). **E:** Time course of inactivation was determined by bi-exponential fits to the inactivating current portion of records obtained using a command potential of +20 mV. This gave fast time constants (τ_f_, 5.90±1.28 ms for WT and 4.21±0.26 ms for C546/7AA) and slow time constants (τ_s_, 70.92±10.29 ms for WT and 48.25±2.60 ms for C546/7AA) shown in Ei and Eii respectively. Values from individual experiments are shown overlain on the mean values. There was no significant difference in the proportion of inactivation current described by τ_f_ (89.7±0.01% for WT and 89.7±0.01 for MT, *P*>0.05, *t*-test; n=9 for WT and n=12 for MT). **F** and G: (F) shows current density plots (mean with data from individual experiments overlain) during atrial action potential (AAP) “AP clamp”. Bar chart shows maximum current density during atrial action potential (AAP) repolarisation for WT (582.81±24.25 pA/pF) and MT (988.73±104.58 pA/pF) Kv4.3S channels (*P*<0.01, *t*-test; n=10 for WT and n=13 for MT). (G) shows representative currents (shown as absolute currents, nA) for WT (black) and MT (red) Kv4.3S current overlaid with an atrial action potential (AAP). Like the normalised (current density) data, absolute current amplitude was increased by the C546/547AA mutations (9.35±0.55 nA for WT and 14.56±1.24 nA for MT, *P*<0.01, *t*-test; n=10 for WT and n=13 for C546/547AA).

An atrial action potential (AP) waveform was applied under voltage clamp (AP clamp) to examine channel behaviour during a dynamic physiological waveform. With an atrial AP command, C546/547AA again showed an increased current amplitude (Figure 5F, 5G). Thus, the gain of function phenotype of the non-palmitoylatable mutant persists with a physiological waveform, consistent with palmitoylation acting as a negative modulator of Kv4.3S current magnitude.

The voltage dependence of channel inactivation was studied with a protocol shown in the upper inset of Figure 6. The V_0.5_ of steady-state inactivation was negatively shifted for non-palmitoylated Kv4.3S compared to WT (Figure 6Ai, 6Aii, 6B, 6C). The time-course of recovery from inactivation was also determined (protocol shown in the lower inset of Figure 6) and the results are illustrated in Figures 6 Di, Dii and E. The time course of recovery from inactivation was fitted with a single-exponential equation to give a recovery time constant (τ_R_). Recovery from inactivation was accelerated for non-palmitoylated Kv4.3S compared to WT (Figure 6E), with a reduced recovery time-constant (τ_rec_, Figure 6F). In summary, S-palmitoylation was observed to limit Kv4.3 magnitude and to have modest but significant effects on the voltage dependence of inactivation and current reactivation.

**Figure 6.**
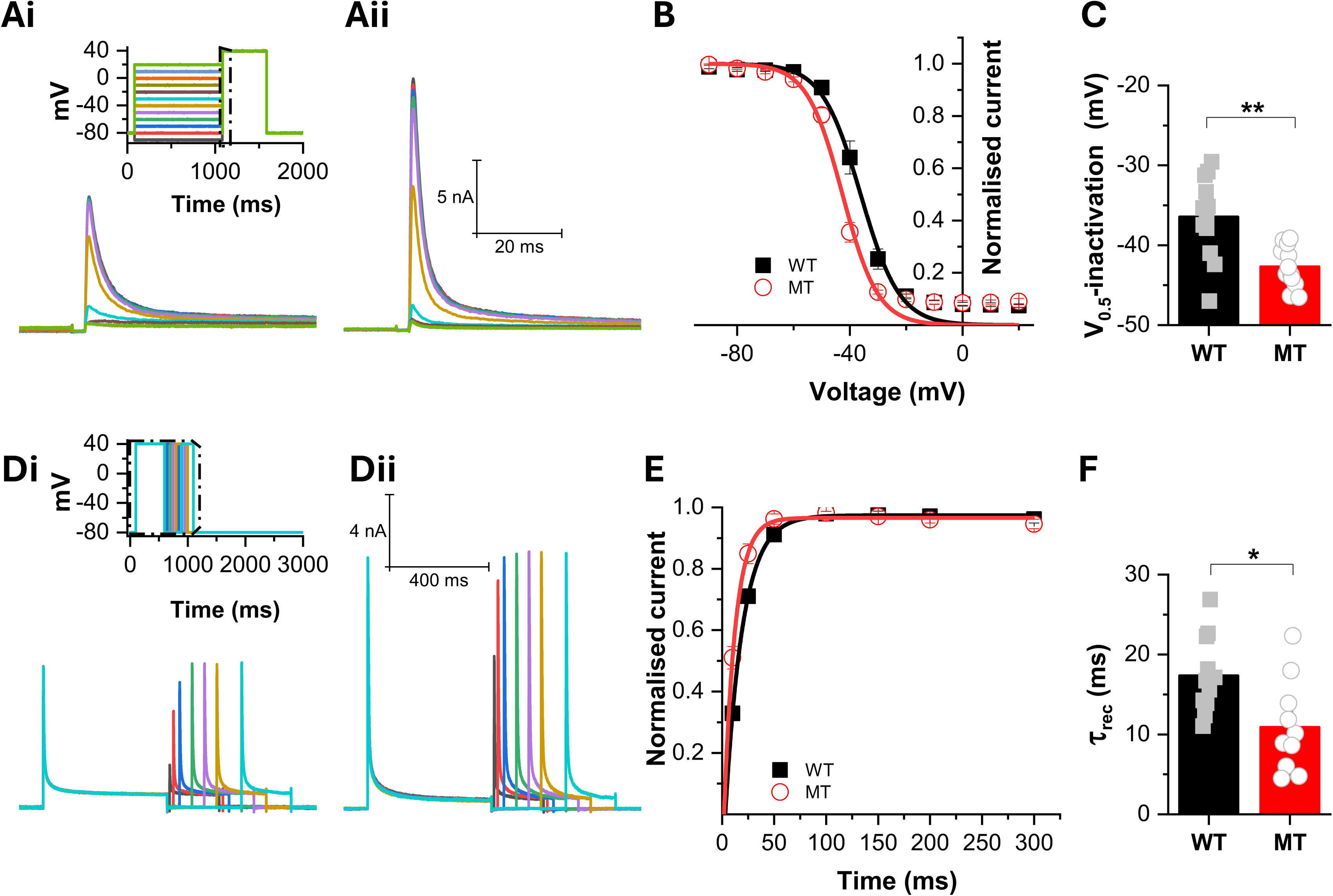
Effects of palmitoylation on Kv4.3S voltage dependence of inactivation and time course of recovery from inactivation. A: Representative families of Kv4.3S current for WT (Ai) and C546/547AA (Aii) channels, elicited by the voltage protocol shown as an inset to Ai to investigate the voltage dependence of inactivation. The traces focus on the current profile upon depolarisation following the conditioning steps (highlighted in the inset by the boxed area). **B** and C: Availability-voltage plots describing Kv4.3S inactivation. The plots were fitted with a Boltzmann equation to derive half-maximal inactivation voltage (V_0.5_, panel C; −36.45±1.48 mV for WT and −42.69±0.75 mV for C546/7AA, *P<*0.01, *t*-test; n= 12 for both WT and MT; values for individual experiments are shown overlain on the mean data plots). There was no significant difference in slope factor (*k*, 5.69±0.22 mV for WT and 5.88±0.38 mV for MT, *P*>0.05, t-test). **D:** Evaluation of recovery from inactivation. Representative current traces of WT (Di) and C546/547AA (Dii) Kv 4.3S channels elicited by the voltage protocol shown as an inset to Di to study the time course of recovery from inactivation. The traces focus on the current profile in the boxed area. **E**,F: Time course of recovery from inactivation. The peak current amplitudes were normalised to the maximal current during the protocol for each cell. The plots were then fitted with one-phase exponential association to give time constant values of recovery (τ_rec_) as shown in panel F (τ_rec_ of 17.36±1.46 ms for WT and 10.89 ±1.84 ms for C546/7AA; *P*<0.05, t-test; n=11 for WT and n=10 for MT), in which results from individual experiments are shown overlain on the mean data.

### Influence of palmitoylation on Kv4.3 protein interactions and function in induced pluripotent stem cell derived cardiomyocytes

The contribution of the intrinsically disordered Kv4.3 C-terminal tail to cardiac I_to_ remains incompletely defined. Multiple post-translational modifications within this region, (including phosphorylation very close to the Kv4.3 palmitoylation sites^56, 57^) are known to modulate Kv4.3 channel properties. SAP-97 (DLG1) binding to the Kv4.3 C terminus stabilises the channel at the plasma membrane,^58^ whereas other elements of the C-terminal tail are destabilising. The conformational flexibility characteristic of intrinsically disordered regions, such as the Kv4.3 C tail, enables context-dependent protein-protein interactions. On this basis, we first tested the hypothesis that palmitoylation of the Kv4.3 C-terminal tail alters its protein interaction landscape in induced pluripotent stem cell-derived cardiomyocytes (hiPSC-CMs). To do this, we fused the proximity-labelling enzyme TurboID to WT or non-palmitoylatable Kv4.3S, expressed these constructs in stem-cell-derived cardiomyocytes, applied a brief biotin pulse to label proximal proteins (Figure 7A), and compared the resulting interactomes using label-free quantitative proteomics (Figure 7B).

**Figure 7:**
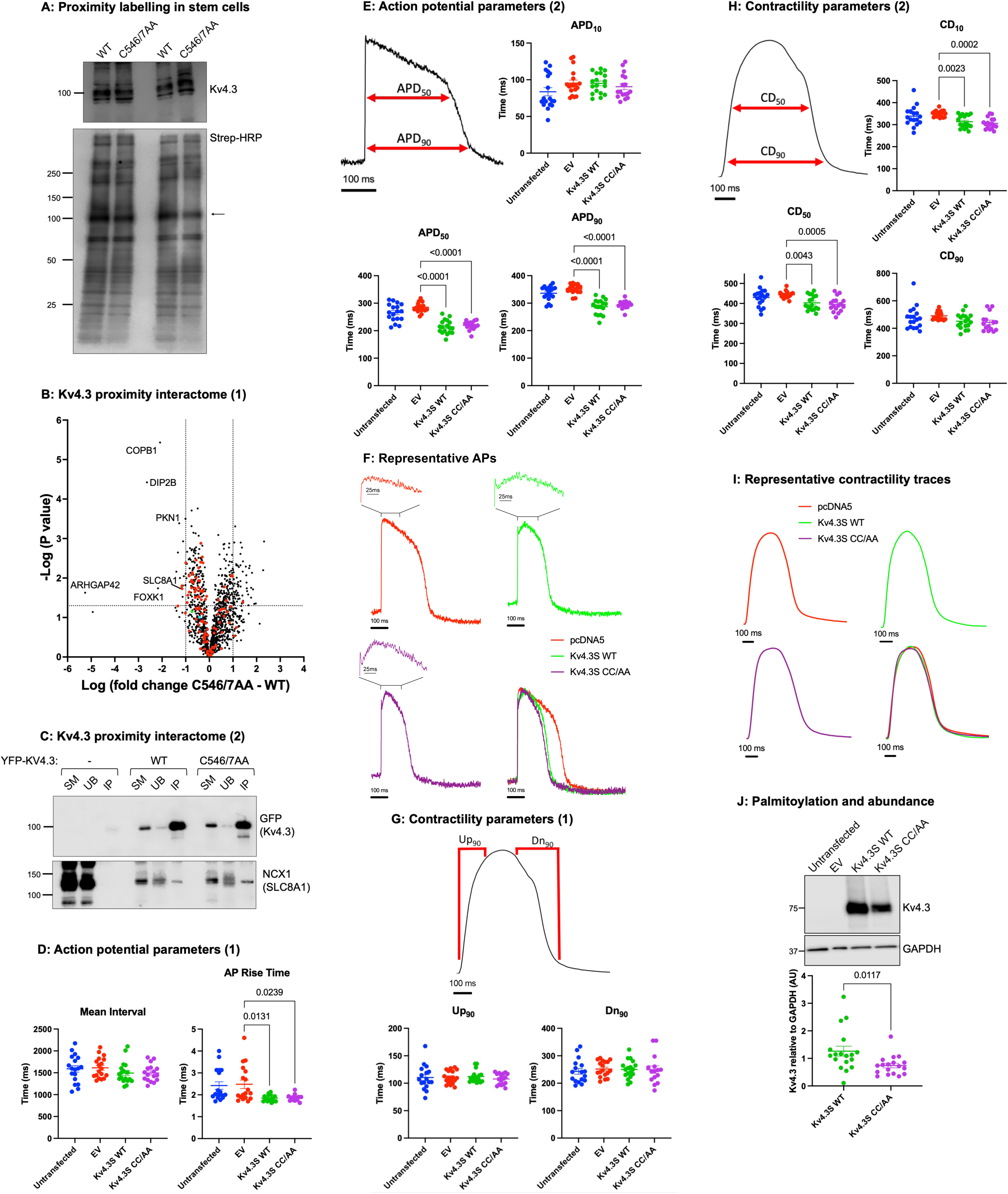
Palmitoylation and Kv4.3 function in stem cell derived cardiomyocytes. A: Kv4.3 proximity labelling in iPSC-CMs. Expression of WT and unpalmitoylatable Kv4.3 fused to TurboID in stem cell derived cardiomyocytes. Upper panel shows expression of the fusion protein. Lower panel shows biotinylated proteins from whole cell lysates detected using streptavidin-HRP. The arrow marks the position of the Kv4.3-TurboID fusion protein biotinylating itself. **B:** Quantitative comparison of Kv4.3 microenvironment remodelling by palmitoylation. Proteins annotated as resident at the cell surface membrane are highlighted in red. Selected proteins are identified. The green dot indicates the bait protein, murine Kv4.3. The cyan dot indicates zDHHC5 (N=4 biological replicates per group). **C:** Interaction of Kv4.3 and NCX1 assessed using co-immunoprecipitation. YFP-Kv4.3 was co-transfected with NCX1 in HEK293 cells and immunoprecipitated. SM: soluble cell lysate, UB: proteins not immunoprecipitated, IP: immunoprecipitated proteins. **D:** Impact of Kv4.3 palmitoylation on AP parameters. iPSC-CMs were transfected with either empty vector (EV), wild type Kv4.3S (WT) or unpalmitoylatable C546/547AA (CC/AA) Kv4.3S, loaded with the voltage sensitive dye FluoVolt, and optical action potential traces recorded using the CellOPTIQ platform. The mean interval between APs was unchanged in all conditions (left: 1590±73 ms, 1613±56 ms, 1494±59 ms, 1495±47 ms for untransfected, EV, Kv4.3S WT, and Kv4.3S CC/AA respectively, P>0.05, ordinary one-way ANOVA). The AP rise time was equally accelerated by expressing WT or CC/AA Kv4.3 (right). Values plotted are mean±SEM, N=13-18 per group over four independent plating days. Outliers were identified using GraphPad Prism 10 outlier test by ROUT (Q=1%) and data analysed with a one-way ANOVA with a Šidák’s post hoc test. **E:** Impact of Kv4.3 palmitoylation on AP kinetics. A representative AP trace annotated with action potential duration at 50% repolarisation (APD_50_) and action potential duration at 90% repolarisation (APD_90_) is shown. APD10 was unchanged and APD50 and APD90 were equally shortened by expressing WT or CC/AA Kv4.3. Values plotted are mean±SEM, N=16-18 per group over four independent plating days. Outliers were identified using GraphPad Prism 10 outlier test by ROUT (Q=1%) and data analysed with a one-way ANOVA with a Šidák’s post hoc test. **F:** Representative AP traces. Example recordings from iPSC-CMs transfected with pcDNA5 (transfection control, upper left), WT Kv4.3S (upper right), CC/AA Kv4.3S (lower left) are shown overlaid (lower right). Zoomed traces of the regions highlighted by the brackets above each AP demonstrate the characteristic spike-and-dome AP morphology indicative of Kv4.3 function. **G:** Impact of Kv4.3 palmitoylation on contraction parameters. iPSC-CMs transfected with either empty vector (EV), wild type Kv4.3S (WT) or unpalmitoylatable C546/547AA (CC/AA) Kv4.3S imaged on the CellOPTIQ platform. A representative contractility trace annotated with Up_90_ (time from 10% to 90% rise of the contractility transient) and Dn_90_ (time from 10% to 90% decay of the contractility transient). Neither parameter was changed by expressing WT or CC/AA Kv4.3. Values plotted are mean±SEM, N=15-18 per group over four independent plating days. **H:** Impact of Kv4.3 palmitoylation on contraction kinetics. A representative contractility trace annotated with CD_50_ (time interval between 50% rise and 50% decay of contraction) and CD_90_ (time interval between 10% rise and 90% decay of contraction) is shown. CD_90_ was unchanged and CD_10_ and CD_50_ were equally shortened by expressing WT or CC/AA Kv4.3. Values plotted are mean±SEM, N=15-18 per group over four independent plating days. Outliers were identified using GraphPad Prism 10 outlier test by ROUT (Q=1%) and data analysed with a one-way ANOVA with a Šidák’s post hoc test. **I:** Representative contractility traces. Example recordings from iPSC-CMs transfected with pcDNA5 (transfection control, upper left), WT Kv4.3S (upper right), CC/AA Kv4.3S (lower left) are shown overlaid (lower right). **J:** Expression of Kv4.3 in hiPSC-CMs. Whole cell lysates from hiPSC-CMs transfected with either empty vector (EV), wild type Kv4.3S (WT) or unpalmitoylatable C546/547AA (CC/AA) Kv4.3S analysed via SDS-PAGE/western blotting for Kv4.3 and GAPDH (loading control). Graph shows expression of Kv4.3 relative to GAPDH, mean±SEM, N=18 per group over four independent plating days, analysed with an unpaired t-test.

We identified a broad Kv4.3 interactome (quantitative data for 1329 proteins, Supplementary Table 2) enriched in abundant cellular proteins. Functional annotation clustering^59^ identified cytoplasmic, cytoskeletal, mitochondrial, and ribosomal components, consistent with the high sensitivity and spatial reach of TurboID-based approaches. A single palmitoylating enzyme, zDHHC5, was detected. Comparative analysis revealed a relatively restricted set of proteins whose proximity to Kv4.3 was altered by S-palmitoylation, suggesting that this modification exerts modest effects on the Kv4.3 microenvironment. Among 103 Kv4.3-proximal proteins annotated as localised at the cell surface membrane (sarcolemma, cell junction, intercalated disk, Figure 7B), the Na/Ca exchanger NCX1 (SLC8A1) is of particular interest. NCX1 is itself regulated by S-palmitoylation and exhibits altered membrane organisation and activity in response to this modification.^42^ The identification of NCX1 enriched within the S-palmitoylated Kv4.3 proximal proteome is therefore consistent with the idea that palmitoylated Kv4.3 resides within specialised membrane microdomains, where it may co-localise with other lipid-regulated transport proteins. NCX1 co-immunoprecipitated with both wild type and non-palmitoylated Kv4.3 (Figure 7C). Hence the overall (and surface-membrane specific) changes in the Kv4.3-proximal proteome were modest, and these data support a model in which palmitoylation regulates the activity of Kv4.3 rather than driving large-scale remodelling of its interaction network.

We next investigated the impact of Kv4.3 S-palmitoylation on electrical activity of hiPSC-CMs by recording APs from transfected monolayers of hiPSC-CMs loaded with the voltage sensitive dye FluoVolt. In hiPSC-CMs, expressing WT Kv4.3S or the non-palmitoylatable C546/547AA mutant did not change the spontaneous beating rate (mean interval) but each shortened AP rise time and AP duration (APD) relative to vector control across APD_50_ and APD_90_ but not APD_10_ (Figure 7D, 7E). There was no difference in AP rise time or duration between WT and C546/547AA, indicating that the expressed Kv4.3 is sufficient to accelerate repolarisation independently of palmitoylation at C546/547 under these conditions. Representative traces show a pronounced phase-1 notch and global APD shortening when Kv4.3S was expressed (Figure 7F). Contractility analysis revealed that expressing WT or non-palmitoylated Kv4.3S did not change contraction or relaxation rates (Up_90_ or Dn_90_, Figure 7G); CD_10_ and CD_50_ were shortened compared to vector control, whereas CD_90_ was unchanged (Figure 7H, 7I). Thus, early relaxation kinetics accelerate with Kv4.3 expression while late relaxation is preserved. Across experiments, the C546/547A mutant accumulated to ∼50% lower steady-state abundance than WT, consistent with reduced expression stability of the non-palmitoylatable form (Figure 7J).

## Discussion

We identify a phosphorylation-dependent mechanism that licenses Kv4.3 S-palmitoylation, and provide a molecular explanation for splice variant specific differences in lipid modification and channel function. Specifically, phosphorylation centred on S538 promotes formation of a palmitoylation-competent Kv4.3 species that depends on intact N-terminal/T1 architecture, is promoted by KChIP2.1, enabling S-palmitoylation at C546/547. In parallel, we define a cis-acting inhibitory element within residues 488-498 of the Kv4.3L splice insert that suppresses palmitoylation and thereby explains the lower S-palmitoylation state of Kv4.3L relative to Kv4.3S.

### Phosphorylation as a priming modification for S-palmitoylation

Our data support a model in which phosphorylation precedes and enables S-palmitoylation of Kv4.3. Multiple independent perturbations, including nanobody-targeted PP2A recruitment, serum withdrawal, and broad kinase inhibition reduced Kv4.3 S-palmitoylation but in some cases only after a delay relative to loss of phosphorylation, indicating temporal uncoupling between a rapidly turning-over phosphorylation event and a slower turning-over lipid modification. This strongly suggests that phosphorylation acts upstream to generate a substrate competent for S-palmitoylation rather than directly stabilising the S-palmitoylated state. The inability of S538D to fully substitute for phosphorylation indicates that authentic phosphate chemistry (rather than simply negative charge) is required for Kv4.3 S-palmitoylation. We suggest that a phospho-specific protein interaction or local hydrogen-bonding geometry, rather than purely electrostatic priming, underlies the requirement.

This mechanism aligns with emerging precedents in cardiac signalling. Most notably, phospholemman phosphorylation at Ser68 enhances its palmitoylation by zDHHC5,^45, 47, 60^ demonstrating that phosphorylation can positively regulate S-palmitoylation by promoting substrate recognition or enzyme engagement. Our findings extend this principle to a voltage-gated potassium channel and suggest that phosphorylation-dependent recruitment to zDHHC enzymes may represent a broader organising principle for dynamic regulation of membrane protein S-palmitoylation.

Consistent with this, proximity labelling, co-immunoprecipitation and modelling collectively support a mechanism in which phosphorylated S538 enhances Kv4.3 interaction with zDHHC5, positioning C546/547 for catalysis. The AlphaFold-guided docking analysis suggests that electrostatic interactions between phosphorylated Kv4.3 residues and positively charged surfaces within the zDHHC5 catalytic loop could stabilise a productive orientation of the intrinsically disordered Kv4.3 C-terminal tail. Importantly, mutation of S538 dramatically reduced Kv4.3 S-palmitoylation, but reduced (and did not abolish) Kv4.3 interaction with zDHHC5, suggesting that phosphorylation helps position Kv4.3 for catalysis rather than creating a binary binding determinant.

Notably, detectable Kv4.3 palmitoylation in zDHHC5-deficient cells indicates that zDHHC5 is not the sole enzyme acting on Kv4.3, and that phosphorylation likely represents a shared upstream priming signal utilised by multiple zDHHC-PATs rather than a strictly enzyme-specific docking event. The larger effect of S538 mutation compared with zDHHC5 ablation further supports this interpretation.

### KChIP2.1 promotes formation of a palmitoylation-competent state

KChIP2.1 increased Kv4.3 palmitoylation without abolishing splice-dependent differences, indicating that it enhances access to the palmitoylation pathway rather than determining substrate specificity. Since palmitoylation levels were normalised to total Kv4.3 abundance, this effect cannot be explained simply by increased Kv4.3 expression. Instead, our data support a model in which KChIP2.1 facilitates formation of a conformation or subcellular context permissive for phosphorylation and subsequent palmitoylation.

The influence of KChIP2.1 on Kv4.3 S-palmitoylation (but not KChIP2.2, which is not S-palmitoylated and therefore cannot localise to membranes^61^) and the dependence on intact N-terminal/T1 architecture suggest that assembly of the Kv4.3-KChIP complex is upstream of phosphorylation and palmitoylation competence. Given that KChIP2 is known to stabilise Kv4 complexes and enhance their surface expression,^61^ one simple interpretation is that complex assembly promotes exposure of the Kv4.3 distal C-tail for modification by membrane-associated kinases and zDHHC enzymes, although direct evidence for trafficking-dependent mechanisms will require further study.

### Functional consequences of palmitoylation

Functionally, palmitoylation constrains Kv4.3 channel activity. Non-palmitoylatable Kv4.3S exhibited increased peak current, accelerated inactivation, left-shifted activation, and faster recovery, indicating that S-palmitoylation modification stabilises channel states associated with reduced availability and slower gating transitions. Phosphorylation sites previously identified in Kv4.3 (including PKC-sensitive T504 and CaMKII-dependent sites) are also located near the Kv4.3 S-palmitoylation sites and are known to regulate current amplitude and inactivation kinetics.^35, 57, 62–64^ Taken together with our data, these findings support a model in which the Kv4.3 C-terminus acts as a regulatory hub integrating multiple PTMs, with phosphorylation and palmitoylation acting cooperatively rather than independently to shape channel behaviour.

Despite these clear effects in heterologous electrophysiology, palmitoylation did not measurably alter AP duration or contractility parameters in hiPSC-derived cardiomyocytes under the conditions tested. Two factors likely contribute. First, expression of Kv4.3 itself is sufficient to accelerate early repolarisation, potentially masking more subtle modulatory effects of palmitoylation. Second, the non-palmitoylatable mutant exhibited reduced steady-state expression, which may partially offset any increase in current magnitude. Together, these data suggest that palmitoylation fine-tunes Kv4.3 function rather than acting as a primary determinant of repolarisation in this model, and that its effects may be more evident under conditions of endogenous expression or pathological remodelling. We do not rule out the possibility that palmitoylation controls steady state expression of Kv4.3 in cardiac cells, but observed no impact of palmitoylation on Kv4.3 stability in engineered cell lines.

### Physiological and pathological implications

In the failing heart, Kv4.3 expression is reduced and splice variant abundance remodels such that Kv4.3L increases and Kv4.3S markedly decreases.^36^ Given that Kv4.3L is less palmitoylated, this shift would be predicted to reduce overall Kv4.3 palmitoylation in heart failure. Since our data indicate that reduced palmitoylation enhances Kv4.3 current amplitude, such remodelling could partially counterbalance the loss of Kv4.3 expression, representing a compensatory mechanism to preserve repolarising current.

More broadly, our findings challenge the conventional view that phosphorylation and S-palmitoylation act antagonistically, instead supporting a model in which rapid phosphorylation events gate access to slower lipid modifications. This hierarchical PTM coupling may be particularly important for ion channels with intrinsically disordered regulatory domains, where conformational flexibility enables long-range communication between modification sites.

### Limitations and future directions

Several limitations should be considered. First, most mechanistic experiments were performed in HEK293 cells, which provide tractability but may not recapitulate cardiomyocyte-specific signalling environments. Second, the kinase responsible for S538 phosphorylation remains unidentified. Our data indicate that it is serum-responsive and staurosporine-sensitive, and the positive impact of KChIP 2.1 but not KChIP2.2 on S-palmitoylation of the TMD-Del construct is consistent with a membrane-associated kinase, but further work is required to define its identity and regulation. Third, reduced expression of certain mutants, particularly within the splice insert, limits interpretation and suggests additional roles for this region in protein stability or quality control. Fourth, although iPSC-derived cardiomyocytes provide a human, genetically tractable model system, their relative structural and electrophysiological immaturity may limit direct extrapolation of our findings to adult ventricular myocardium.

## Conclusion

We define a phosphorylation-dependent PTM cascade controlling Kv4.3 S-palmitoylation, explain splice-variant differences via a discrete inhibitory sequence element, and demonstrate that S-palmitoylation modulates Kv4.3 channel gating. This work establishes Kv4.3 as a model for hierarchical PTM integration, in which phosphorylation primes S-palmitoylation to tune ion channel behaviour.

## Methods

### Ethical Statement

All protocols were approved by the University of Glasgow Animal Welfare and Ethics Review Board.

### Animal Tissue

Hearts and brains from adult male and female mice were collected post-mortem after sacrificing animals using a method designated Schedule 1 by the Animals (Scientific Procedures) Act 1986.

### Plasmids, mutagenesis and cloning

Murine Kv4.3L (cDNA clone MGC:176058 IMAGE:9055709, GenBank: BC141097.1) was obtained from Horizon Discovery. Amino acid numbering refers to the long splice variant throughout. We used In-Fusion HD cloning (Takara Bio) to generate new plasmids and point mutations. Primers were designed using Takara Bio’s online primer design tool. Plasmid DNA (10 ng per reaction) was linearized using forward and reverse primers and CloneAmp Hifi PCR premix (Takara). PCR products were ligated using In-Fusion HD Enzyme premix.

### Cell lines and culture

Human embryonic kidney (HEK) 293 cells were maintained between passage 5-25 in T75 flasks in a humidified incubator at 37°C, 5% CO_2_. Cell lines stably expressing wild-type, mutant and TurboID fusions to Kv4.3 were generated using the Flp-In T-Rex system (Thermo). A HEK-derived zDHHC5 KO cell line was used as previously described.^42, 45^ HEK 293 cells were transfected with plasmid DNA in a 6 or 12 multi-well plate within 24 hours of plating using Lipofectamine 2000 according to manufacturer’s instructions. Cells were harvested 18-20 hours following transfection.

When cells were serum starved, media was supplemented with 50 µM palmitic acid conjugated to 1% BSA to provide a continuous supply of palmitate. For staurosporine treatment, cells were treated either for 1 hour or overnight with 10 µM staurosporine or vehicle (0.1% DMSO).

Human induced pluripotent stem cell-derived cardiomyocytes (hiPSC-CMs) iCell cardiomyocytes^2^ were from Cellular Dynamic International, FUJI Film (catalogue number C1058). The donor background was adult Caucasian female with an apparently normal health status (donor ID 01434, lot number 106448). The tissue source for the cardiomyocytes were fibroblasts that were differentiated following a cardiac differentiation protocol. This results in a mixed population of ventricular, atrial and nodal cells, with the ratio of each cell type subject to variation between batches. HiPSC-CMs were plated and maintained according to manufacturer’s instructions. Cells were transfected 48 hours after plating with one well of a 96-well plate receiving 50 ng of DNA, and 0.2 µl of ViaFect™ (Promega). The following morning, the mixture was removed and 200 µl of fresh media added. AP and contractility recordings were made 48 hours after transfection.

### Resin-assisted capture of acylated proteins (Acyl-RAC) assay to assess S-palmitoylation

S-palmitoylated proteins were purified using resin-assisted capture of acylation proteins (Acyl-RAC).^65^ In brief, cells or pulverised frozen tissues were lysed in blocking buffer (100 mM HEPES, 1 mM EDTA, 2.5% SDS, 1% MMTS) and incubated at 40°C shaking at 1200 RPM for 4 hours to methylate free cysteines. Proteins were precipitated with x3 volumes of 100% acetone at −20°C for 30 minutes minimum to remove excess unreacted MMTS. Pellets were then washed with 70% acetone, dried and redissolved in binding buffer (100 mM HEPES, 1 mM EDTA, 1% SDS). S-palmitoylated proteins were captured by thiopropyl Sepharose beads in the presence of neutral 250 mM hydroxylamine for 2.5 hours with rotation at room temperature. After capture, beads were washed 5x 1 ml with binding buffer. S-palmitoylated proteins were eluted in SDS-loading buffer (100 mM Tris, 4% SDS, 20% glycerol, 0.02% bromophenol blue, 5% β-mercaptoethanol, 100 mM DTT, pH 6.8).

### Co-immunoprecipitation and FLAG tagged proteins

All steps were carried out at 4°C. Anti-FLAG (Merck), anti-HA (Merck), or GFP-TRAP (Proteintech) beads were washed 3×1 ml in lysis buffer (50 mM HEPES pH 7.4, 100 mM NaCl, 50 mM NaF, 5 mM Na_4_P_2_O_4_, 1 mM Na_3_VO_4_, 1 mM EDTA, 1:1000 protease inhibitor cocktail (Merck Millipore), 1% Triton X-100 in dH_2_O) and equilibrated with agitation for 1 hour. Cells were lysed in 500 µl of lysis buffer, insoluble material removed by centrifugation (17,000 G, 5 minutes) and an analytical sample retained representing the whole soluble lysate. The remaining lysate was incubated with equilibrated beads for 2 hours with agitation. The beads were washed 5×1 ml and resuspended in 50 µl 2X SDS-PAGE loading buffer supplemented with 100 mM DTT/ 5% β-ME. Samples were heated (60°C, 10 minutes) and proteins analysed via SDS-PAGE/western blotting.

### Western blotting

In this investigation the following antibodies were used: Kv4.3 (1:200, Alomone APC-017, 1:50 DSHB K75/41), Na pump (1:100, DSHB clone a6F), GAPDH (1:10,000, Sigmaaldrich/MERCK AMAB91152), STREP HRP (1:10,000, Cytiva 10535925), FLAG (1:1000, Sigmaaldrich/MERCK F1804, 1:10,000 Proteintech 80801-2-RR), Flotillin2 (1:2000, BD Biosciences 610383) and GFP (1:5000, Proteintech 3H9). Secondary antibodies used were anti-rabbit conjugated to HRP (Jackson ImmunoResearch, 111-035-144, raised in goat) and anti-mouse conjugated to HRP (Jackson ImmunoResearch, 315-035-003, raised in rabbit).

### Western blot images were acquired using a LI-COR Odyssey FC and bands were quantified using Image Studio Software (LiCOR, version 5.2.5)

To investigate phosphorylation, the Phos-tag^TM^ acrylamide AAL-107 reagent (30 μM) was added to 8% SDS-PAGE gels along with 0.0594 mM manganese chloride to produce a functional Phos-tag ligand. Proteins were separated using 7.5 mA/gel for 100 minutes, followed by 10 mA/gel until the dye front was 0.5 cm from the bottom of the gel. Before transfer, gels were washed 2x 10 minutes in transfer buffer supplemented with 2 mM EDTA.

### Peptide modelling

HPEPDOCK peptide docking programme was used to generate models of zDHHC5 with a phosphorylated Kv4.3 peptide (residues 531-550, including phosphoserine 538).^66^ The receptor co-ordinates came from AlphaFold predicted structure of zDHHC5 (AF-Q9C0B5-F1) and the phosphorylated peptide was created using PyTMs plugin within Pymol.^67^ Default pose prediction settings were used for the docking. The top 20 poses were examined and modelling scenarios in which the Kv4.3 palmitoylated cysteines did not approach the zDHHC5 active site were rejected.

### Single cell electrophysiology

FT293 cells stably expressing wild-type and mutant Kv4.3 channels were passaged with 0.05% trypsin-EDTA (Gibco), gently pelleted and resuspended and seeded onto sterilized glass coverslips in 40 mm Petri-dishes containing fresh medium. Cells were incubated at 37°C (5% CO2) for a minimum of 3 hours following passage prior to any electrophysiological measurements.

Coverslips onto which cells had been plated were mounted on an inverted microscope (Nikon Diaphot) and continuously superfused at 37 ± 1◦C with a modified Tyrode’s solution containing (in mM): 138 NaCl, 4 KCl, 2 CaCl_2_, 1 MgCl_2_, 0.33 NaH_2_PO_4_, 10 Glucose and 10 HEPES (titrated to pH 7.4 with NaOH). Patch-pipettes (Corning 7052 glass, AM Systems Inc.) were pulled (Narishige, PP 830) and polished (Narishige, MF83) to give a final resistance of 2-4 MΩ. The pipette/intracellular solution contained (in mM): 90 KAspartate, 30 KCl, 10 NaCl, 1 MgCl2, 5 EGTA, 5 MgATP, 10 HEPES, 5.5 Glucose (pH adjusted to 7.3 with KOH).^68^ Whole-cell patch-clamp recordings were made with an Axopatch 1D amplifier (Axon instruments) and a CV-4 1/100 head stage. Series resistance values typically lay between 3 and 6 MΩ and were compensated by ∼60-80%. Currents were filtered at 2 kHz and digitised at 10 kHz for all protocols.

### Affinity purification of proteins following TurboID based labelling in FT293 cells

FT293 stably expressing TurboID-Kv4.3 were lysed in 1% Triton X-100, 0.1% SDS, PBS, supplemented with a protease inhibitor cocktail (Merck). Insoluble proteins were removed by centrifugation and biotinylated proteins captured by incubating lysates with streptavidin Sepharose (Cytiva) for 4 h at 4°C. Following extensive washing with 100mM ammonium bicarbonate in 1M urea, captured proteins were reduced and alkylated, beads incubated in trypsin overnight, and peptides prepared for LC-MS/MS analysis as described previously.^54^

### Affinity purification of proteins following TurboID based labelling in stem cell derived cardiomyocytes

Cells were lysed in 1% Triton X-100, 0.1% SDS, PBS, supplemented with a protease inhibitor (PI) cocktail (Merck). Protein concentrations were measured using the DC assay and adjusted to 1 µg/µl in 30 µl of 1% Triton X-100, 0.1% SDS, PBS, PI cocktail. Biotinylated proteins were enriched using an automated KingFisher (KF) Flex system (Thermo Fisher) in KF 96 deep-well plates. 3 µl of Streptadivin magnetic beads (88817, Pierce) were added to each sample in the pull-down (PD) plate and agitated for 2 min at 600 rpm on a plate shaker. The KF PD protocol began with a 2 h mixing step at medium speed, 4 bead collection counts of 20 seconds each, no heating. The beads were subsequently washed with 150 µL RIPA buffer and 150 µL 1% SDS, 50 mM HEPES, 1M urea pH 8.0. Each wash consisted of bead release for 1 min at medium speed, followed by 1 min mixing at medium speed and 4 bead collection counts of 15 s each without heating. Reduction and alkylation were then performed in 150 µl of 0.2% SDS, 10 mM TCEP, 40 mM CAA, 1M urea using the same KF settings as described for the wash steps. The beads were washed twice with 150 µl of 50 mM HEPES, 1 M urea, pH 8.0 using the same settings. In the final step of the PD protocol, the beads were released into 60 µl of 50 mM HEPES, 1 M urea, pH 8.0 for 2 min at slow speed. The KF comb was left in the plate.

### Protein digestion

0.5 μL of 0.2 μg/μL Trypsin in 1M urea/50mM HEPES (V5111, Promega) was added to each sample. Samples were shaken at room temperature for an additional hour with the comb left in the plate to minimise peptide loss from beads remaining on the comb. The plate was then sealed with aluminium adhesive film and incubated overnight at 37 °C with shaking at 650 rpm. The next day the samples were filtered to a protein low bind plate (0030504100, Eppendorf) through the 0.45 µm PVDF filter plates (MSHVN4510, Millipore) by centrifugation at 200 g for 1 min. The filters had been pre-equilibrated with 50 µl of 0.1% formic acid (FA). Following sample filtration, an additional 10 µl of 0.1% FA was passed through the filters into the collection plate to minimise peptide loss. Due to evaporation during digestion, the final sample volume was approximately 50 µl. Sample pH was adjusted to approximately 2-3 by the addition of formic acid (A117-50).

### Evotip loading

Evotips were loaded according to a modified protocol adapted from the manufacturer’s instructions optimized for chemical proteomics.^69^ Briefly, tips were washed with 20 µL of acetonitrile containing 0.1% formic acid, then centrifuged for 1 minute at 800g. Next, they were soaked in propanol for 1 minute, and while still soaking, 20 µL of 0.1% formic acid was applied on top for a second wash, followed by centrifugation at 800g for 1 minute. Half on the sample volume (25 μL) was loaded. After sample loading, the Evotips were washed twice with 125 µL of 0.1% formic acid each time followed by centrifugation for 1 minute at 800g. Finally, 300 µL of 0.1% formic acid was added to each Evotip and spun down for 10 seconds to minimise solvent loss by evaporation.

### LC-MS/MS

Peptides were analysed by nanoLC-MS/MS using an Evosep One (Evosep) coupled with a timsTOF HT (Bruker) equipped with an 8 cm x 150 µm, 1.5 µm analytical column (Evosep). Peptides were separated by the Evosep 60SPD workflow (Analytical solvents A: 0.1% formic acid and B: acetonitrile plus 0.1% formic acid). Column was held at 40°C. Data were acquired in data-dependent acquisition (DDA) PASEF mode with the following settings: m/z range from 100 m/z to 1700 m/z, ion mobility range from 1/K0 = 1.30 to 0.85 Vs/cm2 using equal ion accumulation and ramp times in the dual TIMS analyser of 100 ms each. The collision energy was lowered as a function of increasing ion mobility from 59 eV at 1/K0 = 1.6 Vs/cm2 to 20 eV at 1/K0 = 0.6 Vs/cm2. Isolation width was lowered as a function of decreasing m/s from 3 m/z at 800 m/z to 2 m/z at 700 m/z. Active exclusion was applied for 0.4 min. Each cycle consisted of 4 PASEF ramps (total cycle time 0.53 sec) with 2.75 ms measuring time allowed for each selected precursor. A 50 ng HeLa digest QC sample was run between batches of replicates to verify instrument performance.

### Data processing

ddaPASEF Bruker .d files were processed using Fragpipe 22.0 with the built-in “LFQ MBR” workflow. Briefly, Bruker .d files were searched using MSFragger^70^ (version 4.1) with the following parameters: strict trypsin digestion; up to 2 missed cleavages allowed; precursor ion tolerance: 20 ppm; trimming of protein N-terminal methionine; oxidation (M) and N terminal acetylation as variable modifications; carbamidomethylation (C) as a fixed modification; Human database (Downloaded from UniProt 13 July 2023 containing 20588 proteins with murine Kv4.3-TurboID fasta sequence added) with decoys (50%) and common contaminants added. MSFragger search results were processed using Percolator^71^ (version 3.6.5) for peptide-spectrum match validation, followed by Philosopher^72^ (version 5.1.1) for protein and FDR filtering. Label-free quantification values were calculated using the MaxLFQ algorithm using IonQuant^73^ (version 1.10.27) with match between runs disabled and min ions set as 1.

### Data Analysis

Analysis was carried out in Perseus (v2.1.3.).^74^ Proteins identified only by site, proteins matching the reverse decoy database, and common contaminants were excluded from further analysis. Protein intensity values were log₂-transformed to approximate a normal distribution. Data were subsequently normalised by row-wise mean centring and column-wise median normalisation to reduce technical variation between samples. Proteins were retained for downstream analysis if at least four valid quantitative measurements were present within at least one experimental condition. Missing values were then imputed using random values drawn from a downshifted normal distribution.

### Data Availability Statement

The mass spectrometry proteomics data have been deposited to the ProteomeXchange Consortium via the PRIDE partner repository.

### Action potential and contractility recordings in hiPSC-CMs

HiPSC-CMs were incubated in Tyrode’s solution (composition (in mM): 93 NaCl, 20 NaHCO_3_, 1 Na_2_HPO_4_, 20 sodium acetate, 1 MgSO_4_, 5 KCl, 25 glucose, 1.8 CaCl_2_, 5 HEPES, pH 7.4 at 37°C) for 1 hour and then loaded with 1:1000 FluoVolt (FV) and 1:100 PowerLoad^TM^ concentrate for 20 minutes at 37°C. Tyrode’s-FV solution was then removed and fresh Tyrode’s added. APs were recorded on the CellOPTIQ system (Clyde Biosciences) with an inverted Olympus microscope, 40x objective (0.6NA). Recordings were made at 10 kHz sampling rate for 10 seconds (typically 6-8 APs per recording). Three sites per well were selected for recording. AP recordings were analysed using CellOPTIQ software. The fluorescence traces were baseline subtracted, and the traces then filtered using a 15-pass, 3-point moving average (boxcar) filter. The consecutive APs from a single recording were ensemble-averaged, after aligning timepoints of the first derivative peaks, to generate a single AP trace per recording. Functional measurements reported: mean interval (interval between APs), AP rise time, APD_10_ (time interval between 90% rise and 10% decay of the AP), APD_50_ (time interval between 50% rise and 50% decay of the AP), APD_90_ (time interval between 10% rise and 90% decay of the AP).

Brightfield video recordings of cell contractions were made using an Inverted Olympus microscope with a high-speed Hamamatsu camera, 40x objective (0.6NA), with a recording time of 10 seconds at 100 fps. Cell contractility recordings were analysed using the open source MUSCLEMOTION (MM) contractility algorithm.^75^ This method measures the change in pixel intensity as movement generating a representative contractility trace for each recording. Functional measurements reported: Up_90_ (time interval between the 10% and 90% upstroke levels of contractility), Dn_90_ (time interval between the 90% and 10% levels of the relaxation phase), CD_10_ (time interval between 90% rise and 10% decay of the contraction), CD_50_ (time interval between 50% rise and 50% decay of the contraction), CD_90_ (time interval between 10% rise and 90% decay of the contraction).

### Statistical analysis

Statistical analysis was carried out using GraphPad Prism (Version 10) on a minimum of 3 biological replicates. Data is displayed as mean ± standard error of the mean (SEM) unless stated otherwise. A one-way analysis of variance (ANOVA) with an appropriate post hoc test (Tukey for all groups compared, Dunnett’s for all groups compared to a control and Šidák’s for pre-selected comparisons) was conducted for data sets with more than two groups. For comparisons of two groups, unpaired t-tests were used. Outliers in derived measurements of AP and contractility parameters were identified and excluded using GraphPad Prism (Version 10) outlier test by ROUT (Q=1%). Differences between experimental groups with P-values <0.05 were considered statistically significant.

## Author contributions

Conceptualization JCH, WF

Software FLB

Formal analysis EDM, CD, RJG, SM, OK, JWH

Investigation EDM, CD, CSK, RJG, SM, SA, OK, AW, JWH, DB, JL

Resources ED, FJ, EB

Writing - Original Draft JCH, WF

Writing - Review C Editing All authors

Supervision RCM, GLS, HW, JCH, WF

Funding acquisition EWT, RCM, HW, JCH, WF

---

**Supplementary Table 1** Proteins identified in proximity to TurboID-Kv4.3L in FT293 cells by proximity-dependent biotinylation and LC-MS/MS. The table lists the 691 proteins comprising the Kv4.3 proximal proteome, including the candidate palmitoyl acyltransferase zDHHC5.

**Supplementary Table 2** Quantitative analysis of proteins identified in proximity to wild-type and non-palmitoylatable (C546/547AA) TurboID-Kv4.3S in hiPSC-derived cardiomyocytes by proximity-dependent biotinylation and LC-MS/MS. The table contains 1329 proteins identified as the Kv4.3S proximal proteome, with quantification demonstrating the impact of Kv4.3 S-palmitoylation on its proximal proteome.

